# Maned wolves (Chrysocyon brachyurus) in the Ecological Station of Itirapina-Brazil: temporal and spatial distribution based on long-distance vocalizations recordings

**DOI:** 10.1101/2023.03.03.530946

**Authors:** Angélica Felício da Costa, Linilson Padovese

## Abstract

The maned wolf (Chrysocyon brachyurus; Illiger, 1815) is considered the largest canid in South America, it has crepuscular nocturnal behavior, solitary and skittish habits. Its vocalization has different classifications, one of them called roar-barks, which helps in long-distance communication, being ideal for passive acoustic monitoring. For the study of the vocalizations of C. brachyurus in the Ecological Station of Itirapina in Brazil, were implanted 6 autonomous recorders called EcoPods. The locations of the station with the highest detection of vocalizations, their characteristics, temporal distribution in relation to sunset and moon phases were studied. The point with the highest concentration of roar-barks is the same as the point with the litter’s vocalization, indicating the importance of vocalizations in parental care. Passive acoustic monitoring allows a less invasive study and helps to understand the behavior of the maned wolf for its preservation.

## 1. Introduction

The maned wolf (Chrysocyon brachyurus; Illiger, 1815) is considered the largest canid in South America, and can be found in pastures, shrubby habitats, mixed forest, humid grasslands and savannas. Its presence occurs from the mouth of the Parnaíba River in northeast Brazil, to the south of Paraguay, in the state of *Rio Grande do Sul* and in the west of Peru. There are also records in Argentina and Uruguay (Paula; Dematteo, 2015). Chrysocyon brachyurus is included in the IUCN (International Union for Conservation of Nature) Red List as “vulnerable” (ICMBio, 2018) as it is facing a high risk of extinction in the wild. This fact underscores the importance of monitoring work for the preservation of the species.

Studies of C. brachyurus in captivity (Kleiman, 1972; Rocha, 2011), in nature using a GPS collar (Sabato, et al., 2006; Arrais, 2013; Emmons, 2012), fecal analysis (Aragona; Setz, 2001; Bueno; Belentani; Junior, 2002; Franceschini, 2020) and acoustic monitoring (Ferreira, et al, 2020; Neto, 2021; Dietz, 1984; Rocha, 2011) indicate nocturnal crepuscular behaviors, solitary, monogamous, skittish habits and the importance of vocalizations for interspecific and intraspecific interaction.

The vocalizations of maned wolves have a higher incidence in the months of November to January, due to the breeding season (Brady, 1981). Among the different types of vocalizations, the roar-barks or for Portuguese *Aulido*, is one of the only long-range ones, and ideal for passive acoustic monitoring (MAP) (Ferreira, et al., 2020). The roar-barks are characteristic sounds of C. brachyurus, being similar to those of a domestic dog, however differing by presenting hoarseness, prolongation and repetitions. This type of vocalization has high intensity and low frequency, with a bandwidth of 250 to 1200 Hz. They are most often chanted when the maned wolf is standing, with its head held high and the snout directed upwards (Rocha, 2011).

Even with the relevance of the study of vocalizations for understanding the behavior of maned wolves, there is little research that focuses on the study of their acoustic communication, with works in the 70s and 80s as carried out by Kleiman (1972), Tembrock (1976), Brady (1981) and Dietz (1984) and the most recent by Rocha (2011), Neto (2021), Ferreira, et al., (2020). The MAP allows for a less invasive study of C. brachyuru behavior and enables temporal observation, such as the peak period vocalization in relation to the day, time of year or moon phases; and also the understanding of territorial interaction.

The maned wolf demarcates its area with urine and feces, and may encompass a residence area of 21.7 to 30.0 km^2^ (Dietz, 1984). The preservation of the species is closely related to the protection of the cerrado biome, such as those guaranteed in conservation units (UC) by law n°9.985/2000. The Ecological Station of Itirapina (EEI) is a UC that has one of the last remnants of savanna in the state of São Paulo (ZANCHETTA, et al., 2006) and is home to a maned wolf family.

Through passive acoustic monitoring, the objective of this work is to describe the long-range acoustic communication patterns of C. brachyurus in the Ecological Station of Itirapina from October to December 2021. The raised hypothesis is that there are patterns of vocalization activities such as an increase in evening crepuscular periods and a decrease in these activities on full moon nights. Another hypothesis is that there are points with more and less vocalization frequencies in the Ecological Station. The description of the temporal and spatial profile of vocalizations aims to improve understanding for the preservation of the species.

## 2. Material and Methods

### 2.1. Study Area

The Ecological Station of Itirapina is located in the state of São Paulo, in the municipalities of Itirapina and Brotas, with an area of 2,3000 ha. Its region presents remnants with a predominance of cerrado with savanna and grassland physiognomies. The average annual precipitation is 1,388.7 mm, defining a rainy season from October to March (105 to 311 mm monthly), and a dry season from April to September (18 to 85 mm). As the occurrence of fires is recurrent, there is a large number of native forage species (Silva, et al., 2006). The Itaqueri River and Lobo River Basins are located within the limits of the unit, with small stretches covered with riverside forests that may or may not be floodable; these basins drain the interior of the unit.

In figure 1, which represents the EEI map, it is possible to observe that the surroundings of the conservation unit (UC), the buffer zone, is surrounded by bodies of water and presents anthropic activities. At the bottom of Figure 1, to the south of the map, represented by the gray line, there is a railway line that has constant activity of passing freight train cars. To detect C. brachyurus vocalizations, automatic acoustic recorders were attached to trees at six points in the station.

**Figure 1.**
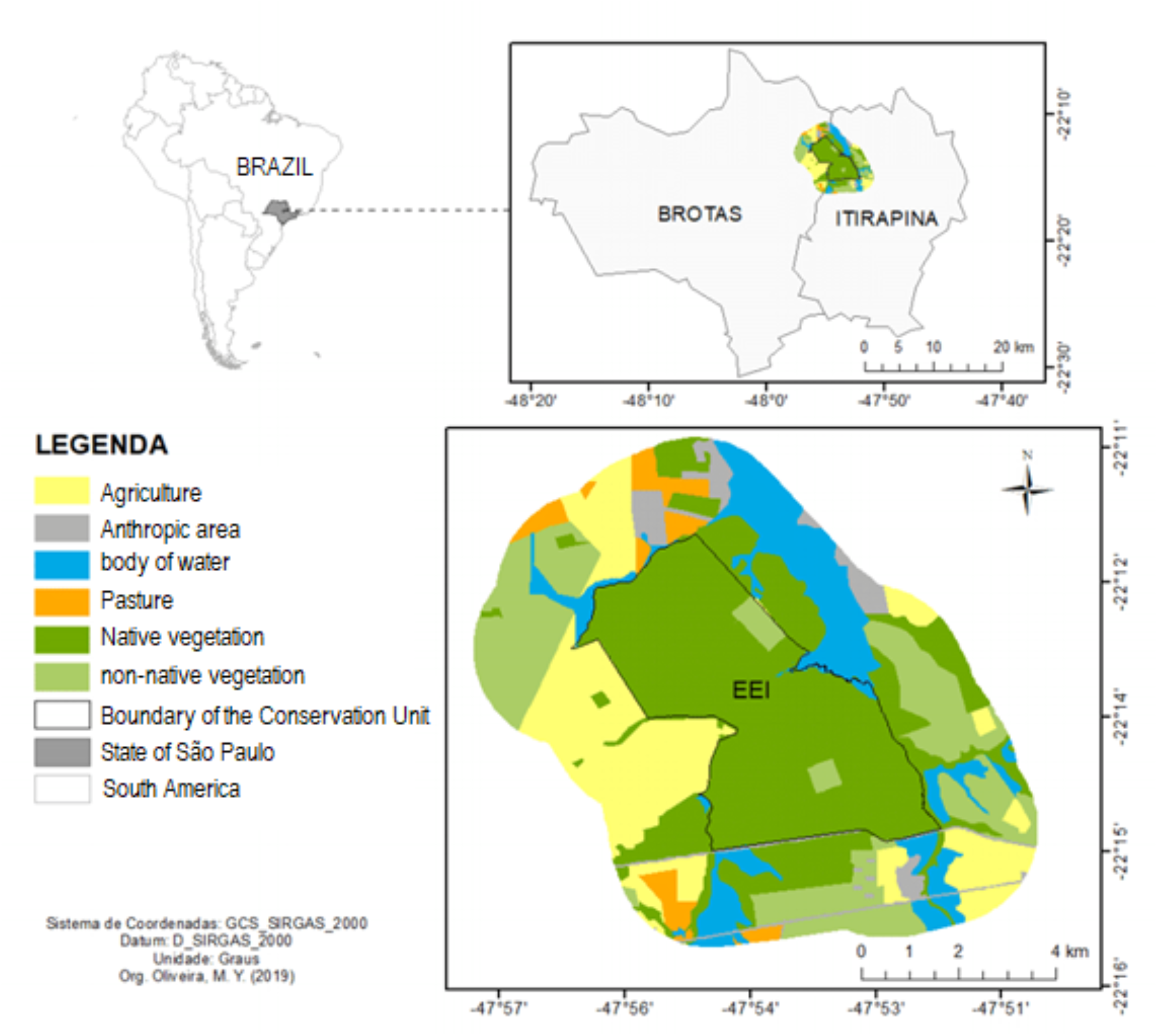
Ecological Station of Itirapina, located between the municipalities of Itirapina and Brotas, in the state of São Paulo. Source: Adapted from Oliveira M. Y. (2019), apud Franceschini (2020).

Were used six autonomous acoustic recorders, called EcoPods, developed by Lacmam (*Laboratório de Acústica e Meio Ambiente da Universidade de São Paulo*-Brazil). The audios were recorded in 16 bits, and 8kHz sampling rate, in 5 minutes files. They were installed at points named A, B, C, D, E and F, as shown on the map in Figure 2.

**Figure 2.**
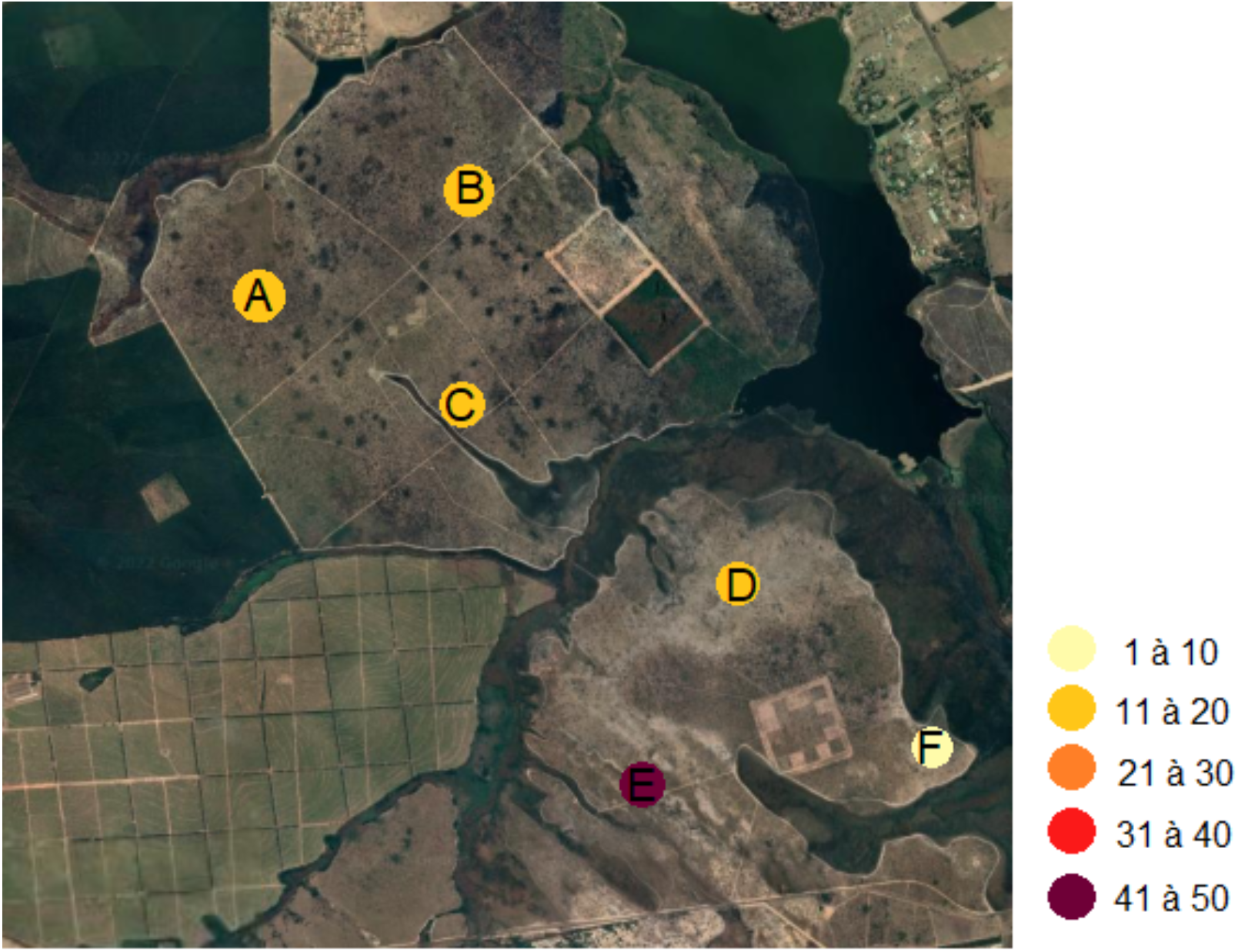
Vocalization sequences of C. brachyurus recorded with six autonomous recorders, EcoPods, at six points indicated by letters A, B, C, D, E and F in the Ecological Station of Itirapina - SP between November and December 2021. The colors represent the approximate amount of vocalization identified per point.

### 2.2. Recordings and Analysis

EcoPods were activated on November 9, 2021, withdrawn on December 24 of the same year and programmed to record for 24 hours. The analyzes and the search for maned wolf vocalizations were performed manually through the spectrograms generated by Raven Pro 1.6 software (Cornell Lab of Ornithology). The search for vocalizations occurred between the period from 5 pm to 5 am due to its peak vocal activity during this interval, as indicated in preliminary studies (Ferreira, et al. 2020; Bueno; Belentani; Junior, 2002; Franceschini, 2020).

Considering a vocalization as a sequence of defined roar-barks and not separated by more than 10 seconds (Ferreira, et al. 2020), the amounts of vocalizations performed by a maned wolf and by more than one were analyzed. The vocalizations performed by one individual were called solo and more than one individual were called interactions. With the survey of the acoustic data of each EcoPod, it was possible to estimate the number of vocalizations from the six points of the Ecological Station of Itirapina.

For the temporal analysis, the vocalizations of each hour after sunset were grouped together and the periods with more and less sound detections of howls, over the 44 nights of recordings were recorded. The relationship between vocalizations and moon phases were also analyzed.

## 3. Results and Discussion

The soundscape of the Ecological Station of Itirapina has registers of anthropophony, such as party music, gunshots and the frequent sound of passing trains. The geophony identified in the recordings was wind and frequent precipitation, due to the period of the rainy season. The biophony detected was biodiverse, composed of different species of birds, insects, anurans and mammals.

### 3.1. Vocalization at Ecological Station of Itirapina

All EcoPods detected roar-barks in the EEI, however there were points with more and others with less vocalization, as shown in figure 2. The points A, B, C and D, which are colored yellow, have detection numbers between 11 and 20. The point F, with a lighter color, had a lower number of vocalizations, only 9. The point E, with a dark purple color, had a higher number of detections of C. brachyurus vocalizations, with 47.

At point A, 13 vocalizations were detected, at point B 13 were also detected; point C had 14 vocalizations. In point D, 16 vocalizations were detected, point F had 9 vocalizations and point E had 45.

The EcoPod located at point E was the only one that captured pup vocalizations. This point is the farthest from human habitation in relation to the others, being close to the Jacaré Guaçu river, which may indicate a favorable environment for parental care.

### 3.2. Solo and interaction vocalizations

Solo vocalizations had higher frequencies compared to vocalization interactive ones at points A, B, D and E, as shown in figure 3. At point C, the same value of solo and interaction vocalizations was detected, and at F interaction detection surpassed the solos.

**Figure 3.**
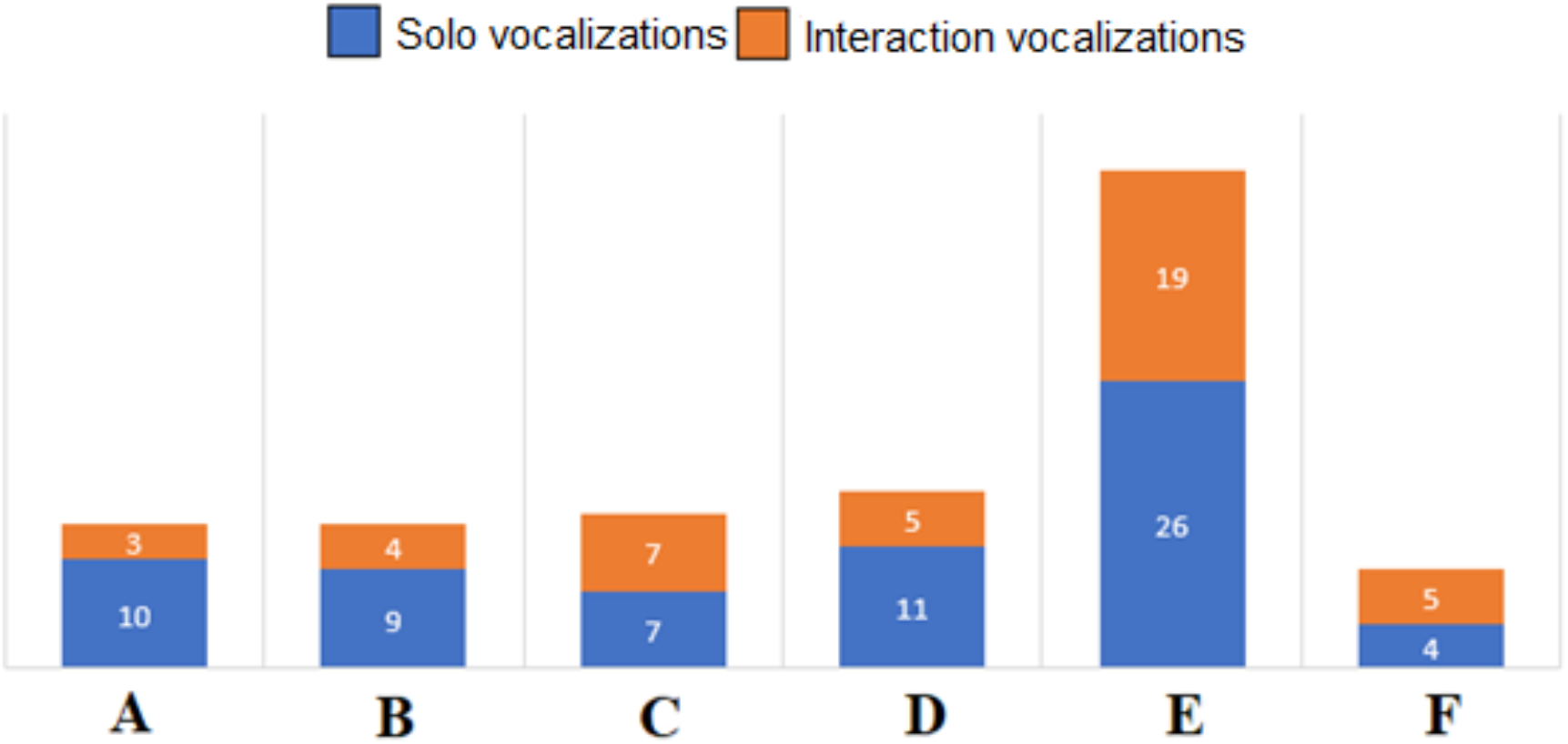
Comparison between solo and interaction vocalizations of *Chrysocyon brachyurus* at points A, B, C, D, E and F located at Ecological Station of Itirapina, SP, Brazil.

The sum of the detections totals 112, with 67 of them being solos; while the interactive ones totaled 40. Therefore, the percentage of solos was 59.8% and of interactions 40.1%. Kleiman (1972) suggests that the interaction between the roar-barks is a spacing and territorial defense mechanism, where an individual answers to another by alternating sequences of roar-barks.

Regarding the vocalization interactions, a high value was observed compared to the work of Ferreira (2020) which presented 17%. The high percentage of interactions could be explained by parental care, since there is detection of baby vocalizations. Acoustic signals help in the parental care of maned wolves, as they establish contact between pups and true parents (Rocha, 2011). The roar-barks help the male to relocate the female who answers it at the place where the offspring are for it to go to them (Neto, 2021; Emmons, et al., 2012).

### 3.3. Vocalizations detected by more than one EcoPod

The same vocalization was detected by more than one EcoPod. This conclusion was possible because they were issued at the same time, had the same number of noises per vocalization and the same duration of time. Of the 112 vocalizations recorded, 31 (27.6%) were detected by two recorders, 5 (4.4%) by three recorders, and 2 (1.2%) by four recorders simultaneously.

### 3.4. Roar-barks by vocalization

Considering a vocalization as a sequence of defined roar-barks not separated by more than 10 seconds, the number of roar-barks per solitary vocalization was counted. Figure 4 exemplifies the spectrogram of a solitary vocalization, each reddish yellow vertical spot represents a roar-barks, in this vocalization there are 13 roar-barks.

**Figure 4.**
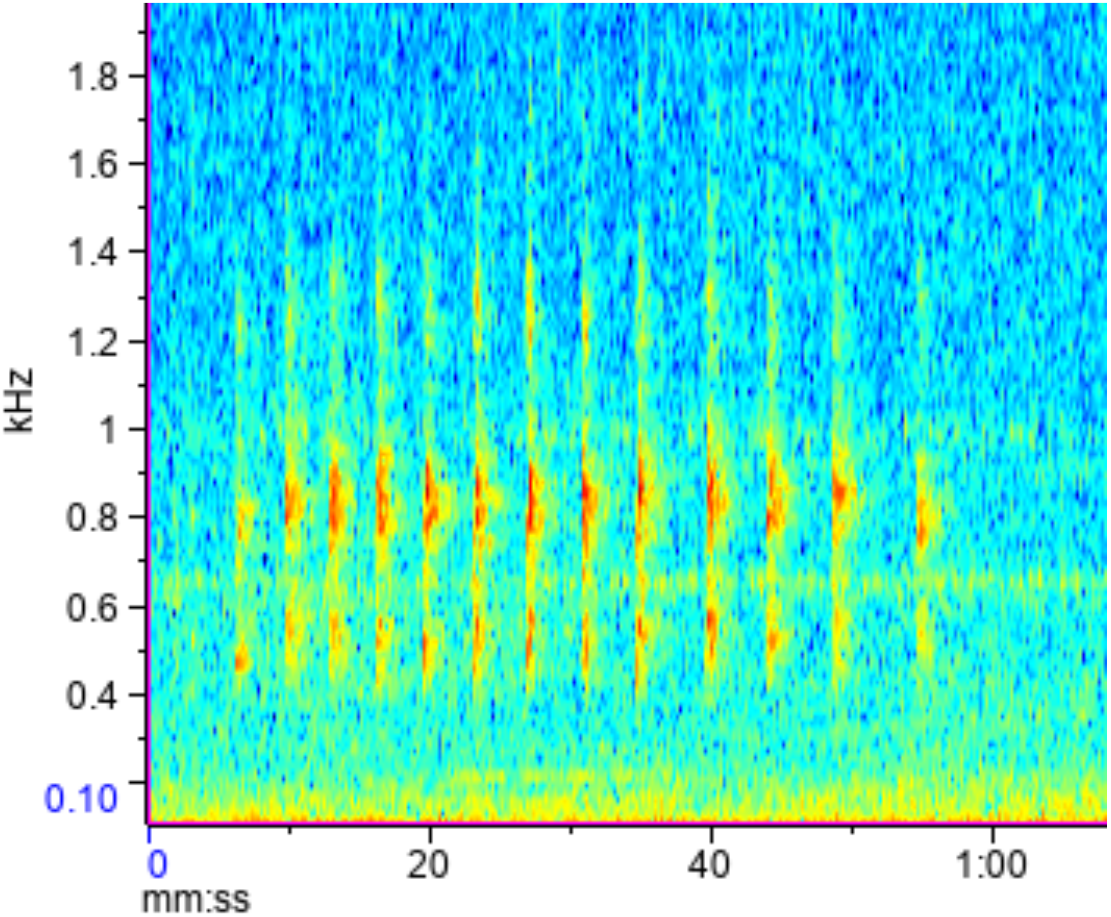
Spectrogram of the vocalization of Chrysocyon brachyurus in the Ecological Station of Itirapina - SP, composed of thirteen roar-barks. The abscissas axis represents the frequency in kilohertz and the ordinates axis the passage of time in minutes.

Through the analysis of 67 solo vocalizations, it was possible to verify that the amount of roar-barks varied from 1 to 30 per vocalization, focusing on values from 5 to 13 as shown in figure 5. The average number of roar-barks per vocalization per sequence was 9.5 (± 5.68).

**Figure 5.**
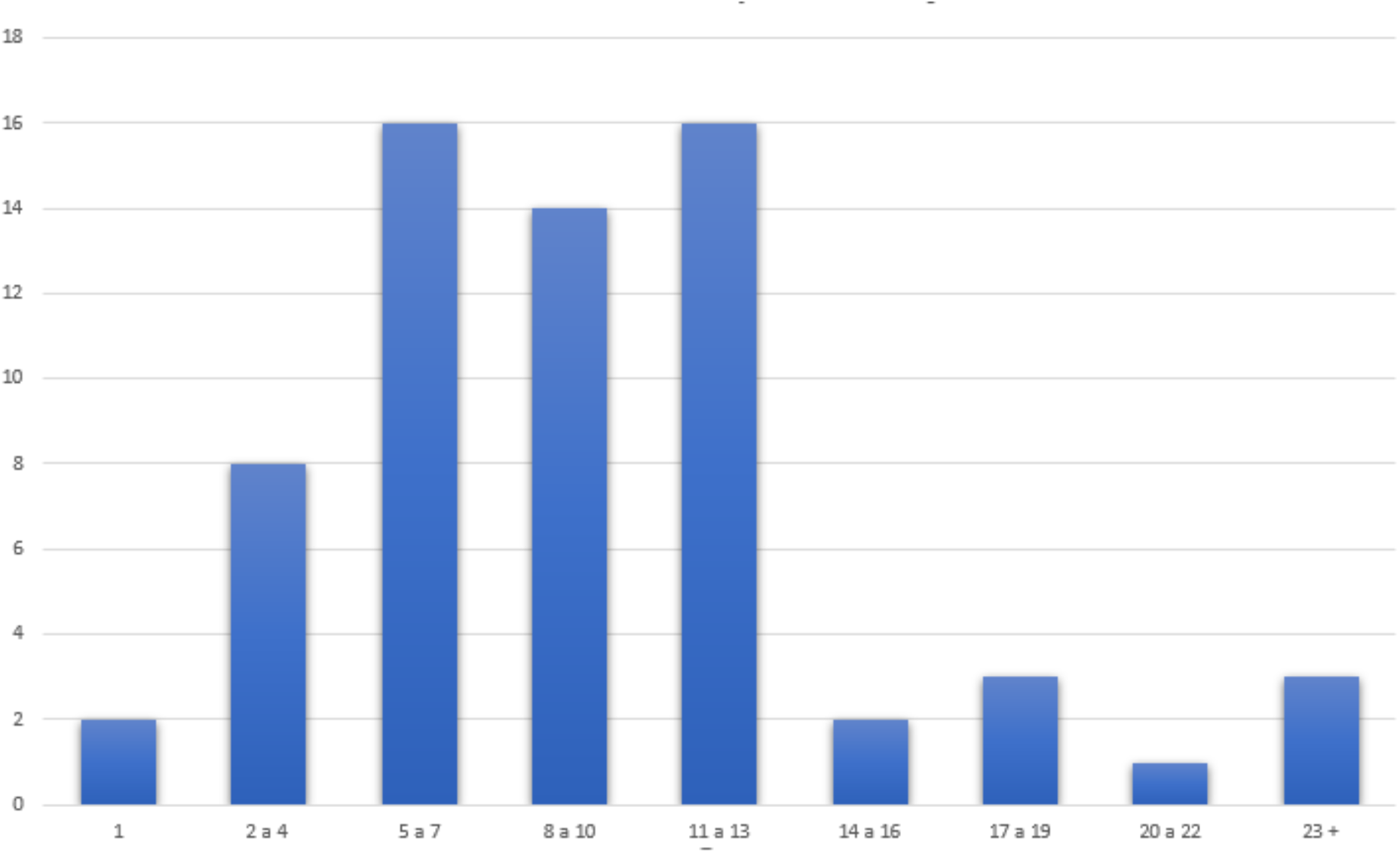
Number of roar-barks sequences in maned wolves’ vocalizations in the Ecological Station of Itirapina - SP. The abscissa axis represents the number of roar-barks per vocalization and the ordinate axis the number of vocalizations. The vocalizations were collected using six autonomous recorders set up at six points in the Station.

### 3.5. Vocalizations after sunset

For this analysis, the same vocalizations captured by more than one recorder were counted as one. For this survey, the hours of each vocalization were cataloged in hours after sunset.

According to the graph in figure 6, it is possible to observe that the period with the highest occurrence of vocalizations was between 5 and 10 hours after sunset, with 73 vocalizations corresponding to 76% of the total detections.

**Figure 6.**
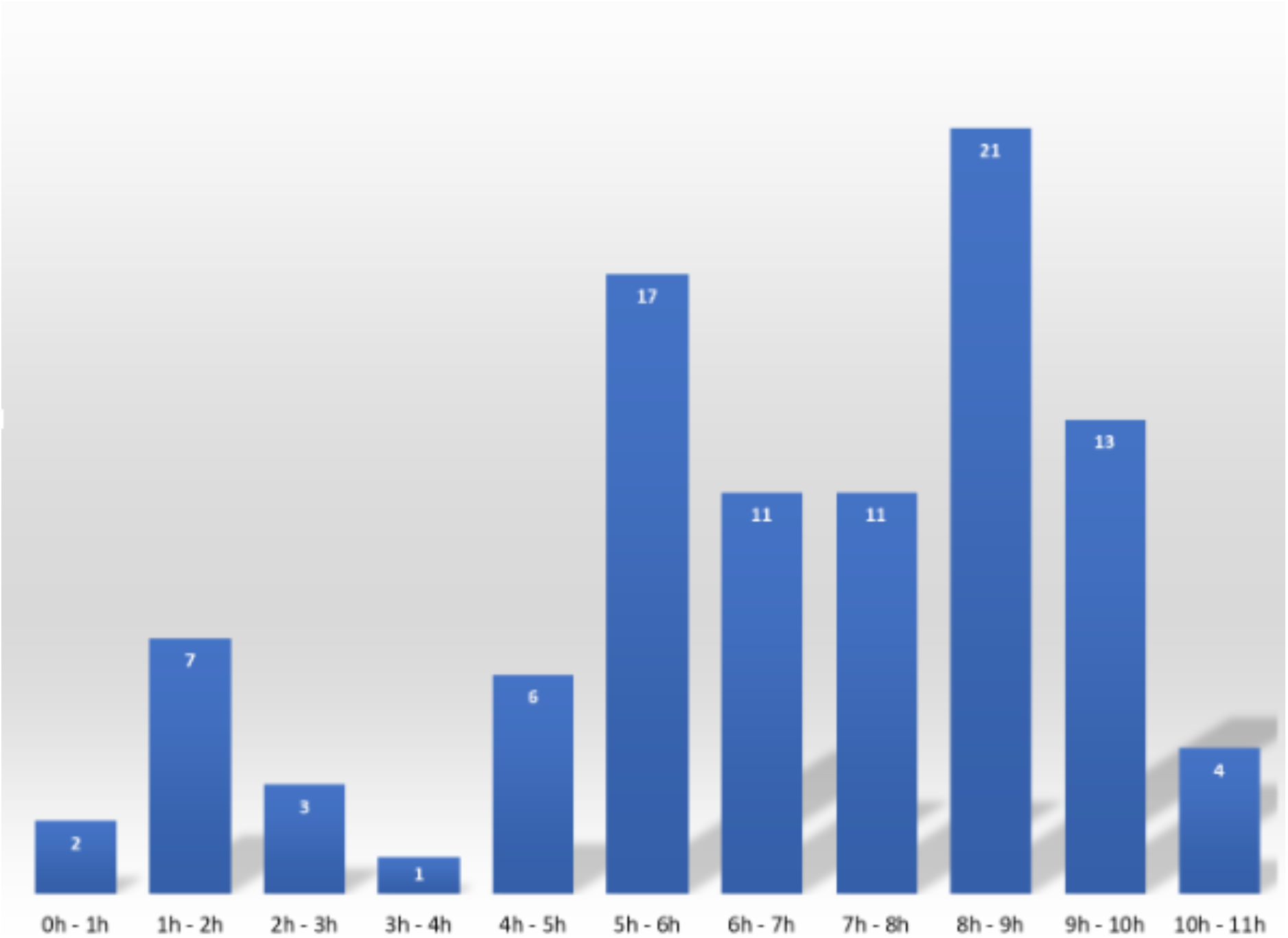
Groupings of numbers of maned wolf vocalization in hours after sunset in the Ecological Station of Itirapina - SP/Brazil.

**Figure 7.**
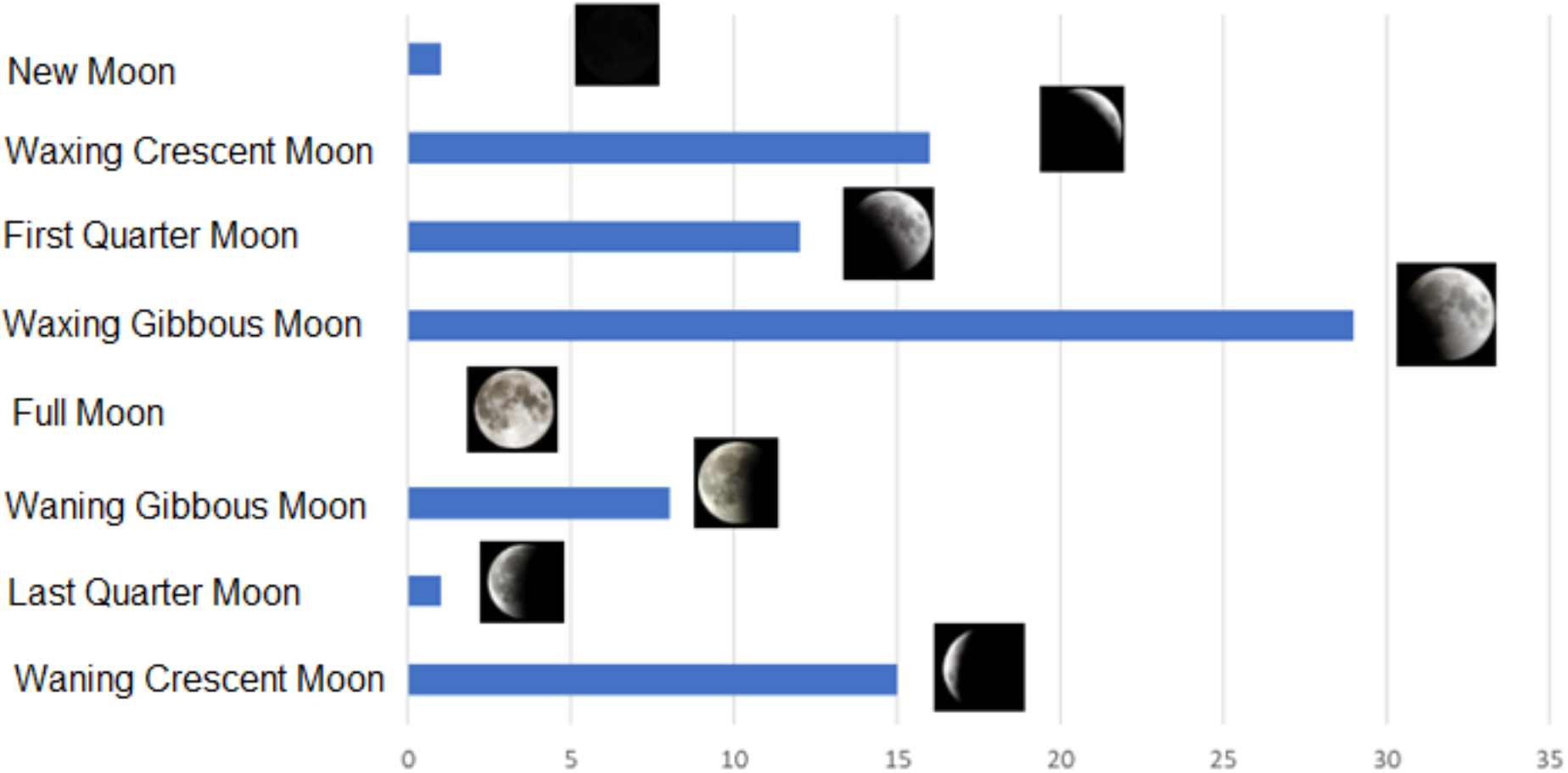
Comparison of the number of vocalizations of Chrysocyon brachyurus with the moon phases, being new moon, waxing crescent moon, first quarter moon, waxing gibbous moon, full moon, waning gibbous moon, last quarter moon and waning crescent moon.

In a general analysis, it is observed that within six hours after sunset there is low detection of vocalizations. Within 2 hours of sunset, only 9.3% were detected and between 2 and 6 hours, 4.1% of the total were detected. After 4 hours there is a significant increase in vocalizations, as between 4 and 6 am there were 23.9%; between 6 and 8 hours 22.9%; between 8 and 10 hours 35% of vocalizations. After 10 hours of sunset the amount of vocalizations decreased to 4.1%.

In the work by Ferreira, et al., (2020), in *Serra da Canastra*, it was observed that the vocal activity of C. brachyurus was concentrated in the first hours after sunset, different from what was observed in this work, which presented a low percentage of vocalizations in the first hours of dusk, which rejects the initial hypothesis.

### 3.6 Vocalization and moon phases

The comparison between moon phases and the vocalizations, represented in image 7, shows that the highest number of vocalizations occurred on a waxing gibbous moon night (with a total of 29 vocalizations), followed by a waxing crescent moon (16 vocalizations), waning crescent moon (15 vocalizations), first quarter moon (12 vocalizations), waxing gibbous moon (8 vocalizations), last quarter moon (1 vocalizations), new moon (1 vocalization). There was no record of vocalization over the forty-four nights of recording during the full moon phase. The work by Ferreira, et al., (2020) also observed a higher concentration of C. brachyurus vocalization in the crescent gibbous phase and low occurrence in the night of full moon and new moon, indicating that the moon phases with moderate lighting increase the frequency of vocalizations, mainly in the phase closest to the full moon.

Even with a higher frequency of vocalizations during the waxing gibbous moon phase, there are no records of vocalizations on full moon nights, which can be explained by the low activity and displacement of the maned wolf on clearer nights (Sabato et al. 2006). On full moon nights, there is a reduction in the activity of prey species, due to the ease of being seen and predated, and therefore, periods with low energy productivity for C. brachyurus.

## 4. Conclusions

Passive acoustic monitoring allowed a less invasive study of maned wolf behavior, enabling temporal and territorial observation. Point E, located southwest of Ecological Station of Itirapina, Brazil, showed a higher detection of vocalizations and the presence of offspring, indicating the importance of vocal communication in parental care. Regarding the temporal analysis, a higher number of vocalizations were detected after 8 to 9 hours of sunset, rejecting the initial hypothesis that there would be more vocalizations in the first hours of evening twilight. Nights of the gibbous crescent moon phase, which precedes the full moon nights, showed higher frequencies of vocal activities, while on the night of the full moon no vocalizations were detected, being periods that require low energy expenditure, mainly for high-energy vocalizations, like the roar-barks.

## References

Aragona, M., Setz, E. Z. F. Diet of the maned wolf, Chrysocyon brachyurus (Mammalia: Canidae), during wet and dry seasons at Ibitipoca State Park, Brazil. Journal of Zoology, v. 254, n. 1, p.131–136, 2001.

Arrais, R. C. Ocorrência de patógenos transmitidos por carrapatos (Anaplasma spp, Babesia spp, Ehrlichia spp, Hepatozoon spp e Rickettsia spp) em lobos guará (Chrysocyon brachyurus) e cães domésticos na região do Parque Nacional da Serra da Canastra, Minas Gerais, Brasil. 2013. Tese de Doutorado.Universidade de São Paulo.

Brady, C. A. The vocal repertoires of the bush dog (Speothos venaticus), crab-eating fox (Cerdocyon thous), and maned wolf (Chrysocyon brachyurus). Animal Behaviour, v. 29, n. 3, p. 649–669, 1981.

Bueno, A. A., Belentani, S. C. S., Motta-Junior, J. C. Feeding ecology of the maned wolf, Chrysocyon brachyurus (Illiger, 1815)(Mammalia: Canidae), in the Ecological Stationof Itirapina, São Paulo State, Brazil. Biota Neotropica, v. 2, p. 1–9, 2002.

Dietz, J. M. Ecology and social organization of the maned wolf (Chrysocyon brachyurus). 1984.

Emmons, L. H. The maned wolves of Noel Kempff Mercado National Park. Smithsonian Contributions to Zoology, 2012.

Ferreira, L. S. et al. Temporal and spatial patterns of the long-range calls of maned wolves (Chrysocyon brachyurus). Mastozoología neotropical, v. 27, n. 1, p. 81–95, 2020.

Franceschini, J. V. Influência da paisagem na dieta de lobo-guará (Chrysocyon brachyurus, Illiger 1815). 2020.

Kleiman, D. G. Social behavior of the maned wolf (Chrysocyon brachyurus) and bush dog (Speothos venaticus): a study in contrast. Journal of Mammalogy, v. 53, n. 4, p. 791–806, 1972.

Ministério do Meio Ambiente. Lista Nacional Oficial de espécies da fauna ameaçadas de extinção. Diário Oficial da União. Brasil, 2018.

Neto, E. T. N. Comunicação acústica de longa distância de lobos-guará de vida livre ao longo das estações reprodutiva e não reprodutiva. 2021. Trabalho de Conclusão de Curso. Universidade Federal do Rio Grande do Norte.

Paula, R. C., Dematteo, K. Chrysocyon brachyurus. The lUCN Red List of Threatened Species 2015: e. T4819A88135664. 2015.

Rocha, V. S. Aspectos do comportamento acústico do lobo-guará chrysocyon brachyurus (Illiger 1815). 2011.

Sábato, M. A. L., Melo, L. F. B., Magni, E. M.V., Young, R. J., Coelho, C. M. A note on the effect of the full moon on the activity of wild maned wolves, Chrysocyon brachyurus. Behavioural processes, v. 73, n. 2, p. 228–230, 2006.

Tembrock, G. Canid vocalizations. Behavioural Processes, v. 1, n. 1, p. 57–75, 1976.

Zanchetta, D., Silva, C. E. F., Reis, C. M., Silva, D. A., Luca, E. D., Fernandes, F. D. S., … & Sawaya, R. (2006). Plano de manejo integrado-Estações Ecológica e Experimental de Itirapina. Instituto Florestal, São Paulo CAPÍTULO, 1.

